# TLR4-mediated neuroimmune signalling drives proprioceptive neuron degeneration in Friedreich ataxia

**DOI:** 10.64898/2026.04.28.721289

**Authors:** Deepika M Chellapandi, Peio A Uhart, Federica Pilotto, Hélène Puccio

## Abstract

Friedreich ataxia (FA) is one of the most common inherited autosomal recessive neurodegenerative disorder, caused by a GAA repeat expansion in the *FXN* gene, leading to frataxin deficiency and progressive sensory and spinocerebellar ataxia. FA is characterized by selective vulnerability of proprioceptive sensory neurons (pSNs) in the dorsal root ganglia (DRG), which are major contributors to sensory ataxia. Despite the central role of mitochondrial dysfunction in FA, the mechanisms that drives the neuronal loss remains unclear. Here, we show that metabolic stress in sensory neurons elicits a neuroimmune response in surrounding tissue, revealing a non-cell-autonomous mechanism of disease progression. We identify Toll-like receptor 4 (TLR4) signaling as a key link between neuronal dysfunction and inflammatory response. Inhibition of TLR4 reduces cellular stress, restores neuronal integrity, and delays disease progression *in vivo*. These findings redefine FA as a disorder involving neuroimmune crosstalk and highlight TLR4 signaling as a potential therapeutic target.

## Introduction

Friedreich ataxia (FA) is the most common inherited ataxia and is characterized by progressive sensory and spinocerebellar ataxia, often accompanied by cardiomyopathy and metabolic dysfunction [1, 2]. The disease is caused by a GAA trinucleotide repeat expansion in the first intron of the *FXN* gene, leading to transcriptional repression and reduced levels of frataxin, a mitochondrial protein essential for iron–sulfur (Fe-S) cluster biogenesis [1, 3]. Frataxin deficiency results in mitochondrial dysfunction, impaired energy metabolism, and increased oxidative stress, which are central features of FA pathophysiology[4, 5].

One of the earliest pathological features of FA is the degeneration of large proprioceptive sensory neurons (pSNs) located in the dorsal root ganglia (DRG). Loss of these neurons disrupts proprioceptive circuits and contributes to sensory ataxia [6, 7]. Despite the well-established role of mitochondrial dysfunction in FA, the mechanisms underlying the selective vulnerability and degeneration of proprioceptive neurons remain incompletely understood.

The DRG represents a complex cellular microenvironment composed not only of sensory neurons but also of satellite glial cells (SGCs), macrophages, Schwann cells, and other stromal populations [8–11]. Increasing evidence indicates that interactions between these cell types play critical roles in maintaining neuronal homeostasis and shaping responses to injury and disease [8, 9, 11, 12]. In particular, SGCs and immune cells have been implicated in neuroinflammatory processes associated with peripheral neuropathies [11, 13–15], raising the possibility that non-cell-autonomous mechanisms contribute to neuronal dysfunction.

Although neuroinflammatory responses have been reported in FA, including observations of satellite cell alterations and inflammatory features in patient DRGs, they remained incompletely characterized in both central and peripheral nervous systems. In particular, how neuron-intrinsic metabolic stress induced by frataxin deficiency influences surrounding glial and immune populations is not well understood.

To address this question, we used the *Pvalb*-Cre conditional frataxin knockout (*Pvalb*-cKO) mouse model, in which frataxin deletion is restricted to parvalbumin-expressing neurons, including pSNs. This model recapitulates key features of FA neuropathology and provides a unique system to investigate how neuron-specific defects impact the surrounding cellular environment. We combined single-cell RNA sequencing with molecular, cellular and *in vivo* approaches to characterize cell-type–specific responses within the DRG and to identify pathways linking neuronal dysfunction to microenvironment changes. Our findings uncover a non-cell-autonomous neuroimmune mechanism linking neuron-intrinsic metabolic stress to inflammatory signaling within the DRG microenvironment and identify TLR4 as a potential therapeutic target in FA.

## Results

### Single-cell RNA sequencing reveals cell-type-specific transcriptional alterations in *Pvalb-* cKO Mice

To identify dysregulated pathways associated with frataxin deficiency in pSNs, we performed single-cell RNA sequencing (scRNA-seq) on DRG isolated from *Pvalb*-cKO mice, in which frataxin deletion is restricted to the proprioceptive sensory neurons pSNs [16]. Lumbar DRGs (L1-L5) were collected at pre-symptomatic (3.5 weeks) and post-symptomatic (7.5 weeks) stages. Following dissociation, approximately 3,000-5,000 cells per sample were sequenced with high-quality transcript capture (Figure S1A). After quality control filtering, the data were normalized and scaled, and unsupervised clustering was performed to identify distinct cellular populations. This analysis identified ten clusters, nine of which were annotated based on canonical marker expression, while one cluster remains unassigned (Figure 1A; S1B). Major DRG cell types were represented, including sensory neurons, satellite glial cells, macrophages, Schwann cells, connective tissue, and pericytes (Figure 1A).

**Figure 1:**
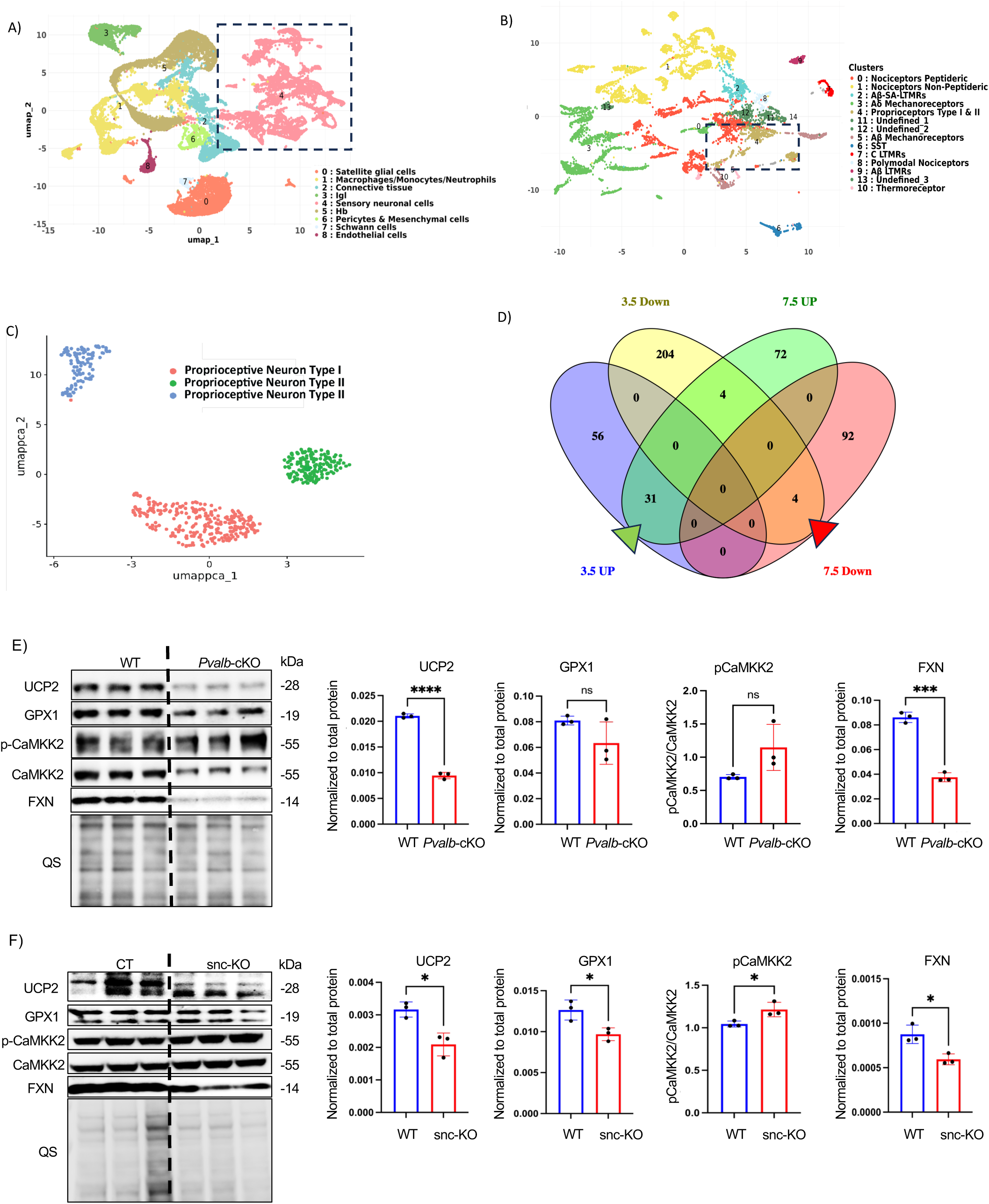
Single-cell transcriptomic profiling of lumbar DRGs from *Pvalb*-cKO mice reveals proprioceptive neuron-specific transcriptional alterations. **(A)** UMAP projection of integrated scRNA-seq data from lumbar (L1–L5) dorsal root ganglia (DRGs) of control and *Pvalb*-cKO mice. Unsupervised clustering combined with canonical marker analysis identified major DRG cell populations, including sensory neurons, satellite glial cells, macrophages, monocytes, Schwann cells, endothelial cells, connective tissue cells, pericytes/mesenchymal cells, and an unidentified cluster. **(B)** UMAP visualization of re-clustered sensory neurons, resolving distinct neuronal subtypes including proprioceptive neuron type I and type II populations, nociceptors, mechanoreceptors, and thermoreceptors. **(C)** UMAP representation of isolated proprioceptive sensory neurons (pSNs), segregating into three transcriptionally distinct clusters corresponding to type I and type II subtypes. **(D)** Venn diagram showing overlap of differentially expressed genes in pSNs at pre-symptomatic (3.5 weeks) and post-symptomatic (7.5 weeks) stages. Upregulated and downregulated gene sets are indicated. (**E)** Immunoblot analysis of selected proteins in whole DRG lysates from post-symptomatic *Pvalb*-cKO mice. Representative blots and quantification of UCP2, GPX1, phosphorylated CaMKK2 (p-CaMKK2), total CaMKK2, and frataxin (FXN) are shown. Protein levels were normalized to total protein (QS loading control). Data are presented as mean ± SEM; statistical analysis was performed using unpaired two-tailed Student’s t-test. ns, not significant; ***P = 0.0001, ****P < 0.0001.**(F)** Immunoblot analysis of primary sensory neuron cultures following frataxin depletion (snc-KO) compared to wild-type (CT) controls. Representative blots and quantification of UCP2, GPX1, p-CaMKK2, total CaMKK2, and FXN are shown. Protein levels were normalized to total protein and expressed relative to control. Data represent mean ± SEM from independent biological replicates (n=3); *P < 0.05.

To increase neuronal subtype resolution, sensory neurons were subset and re-clustered, yielding fourteen distinct clusters. Ten clusters were annotated as defined neuronal types and subtypes, whereas four remained unassigned (Figure 1B; S1C). Given that frataxin deletion is restricted to pSNs in this model, we further isolated pSN populations for focused analysis, resulting in 70–350 cells per sample (Figure S1D). Re-clustering of pSNs identified three transcriptionally distinct clusters, corresponding to one type I and two type II pSN populations (Figure 1C). The two type II clusters likely represent type IIa and IIb subtypes, which could not be definitively distinguished based on transcriptional signature.

### Identification of dysregulated pathways in proprioceptive neurons following frataxin depletion

Differential expression analysis on frataxin-deficient pSNs identified 304 dysregulated genes at pre-symptomatic stage and 200 genes at the post-symptomatic stage. The reduced number of differentially expressed genes at the post-symptomatic stage likely reflects the lower number pSNs recovered in these samples (Figure S1D). As frataxin deletion is restricted to proprioceptive sensory neurons, these data enable the identification of neuron-intrinsic transcriptional alteration. Pathway enrichment analysis revealed dysregulation of canonical FA-associated pathways at both disease stages. KEGG analysis of pre-symptomatic pSNs showed significant enrichment of pathways related to mitochondrial function, oxidative phosphorylation, and neurodegenerative disease signatures. In contrast, post-symptomatic samples exhibited enrichment of signaling pathways associated with neuronal dysfunction and cellular stress response (Figure S2A; S2C). Consistently, Gene Ontology (GO) biological process analysis identified alterations in ion transport, cellular metabolism, and stress-related pathways at the pre-symptomatic stage, with a shift toward dysregulation of neurodevelopmental, migratory, and differentiation-associated processes at the post-symptomatic stage (Figure S2B; S2D).

To identify shared molecular alterations across disease stages, we next focused on genes commonly dysregulated at both time points. Among these, *Gpx1* and *Ucp2* were consistently downregulated (Figure S2E,F)., while *Hsp90aa1*, *Hsp90ab1*, and *Camkk2* were significantly upregulated (Figure S2E,F). This expression profile is consistent with altered redox regulation and activation of stress-responsive signaling pathways in frataxin-deficient proprioceptive neurons [1–6].

### Validation of the scRNAseq finding in DRG tissue and primary sensory neurons

To determine whether transcriptional alterations identified by scRNA-seq were reflected at the protein level, we performed immunoblot analysis on lumbar (L1-L5) DRG lysate from *Pvalb*-cKO mice at pre- and post-symptomatic stages. Proteins were selected based on their consistent dysregulation across disease stages and their functional association with mitochondrial regulation and oxidative stress [22–24, 24–29].

At the pre-symptomatic stage, immunoblot analysis did not detect significant differences in UCP2, GPX1, or GPX4 protein levels between control and *Pvalb*-cKO mice (data not shown). At the post-symptomatic stage, protein levels remained largely unchanged, with the exception of frataxin depletion and a significant reduction in UCP2 (Figure 1E). Given that frataxin deletion is restricted to pSNs, which represent a small fraction of total DRG cells, whole-tissue lysates may dilute neuron-specific protein changes. To address this, we examined primary sensory neuron cultures derived from conditional *Fxn* mice, in which frataxin was selectively deleted *in vitro* [30]. In this enriched neuronal system, UCP2 and GPX1 protein levels were significantly reduced in FXN-deficient neurons (snc-KO) compared to controls (approximately 25% decrease; Figure 1F). In parallel, the ratio of phosphorylated CamKK2 to total CAMKK2 was significantly increased in the frataxin-deficient neurons (Figure 1F).

### TLR4 signaling is activated in frataxin-deficient sensory neurons and dorsal root ganglia

Following validation of transcriptional alterations, we sought to identify upstream regulatory pathways linking these molecular changes. Several of the dysregulated genes identified are functionally associated with Toll-like receptor 4 (TLR4), suggesting this pathway as a potential integrator of neuronal stress response [17, 20, 21, 31–34]. Given the central role of TLR4 in stress and inflammatory signaling pathways[35–37] (Figure 2A), we investigated whether TLR4 signaling is activated in frataxin-deficient sensory neurons.

**Figure 2:**
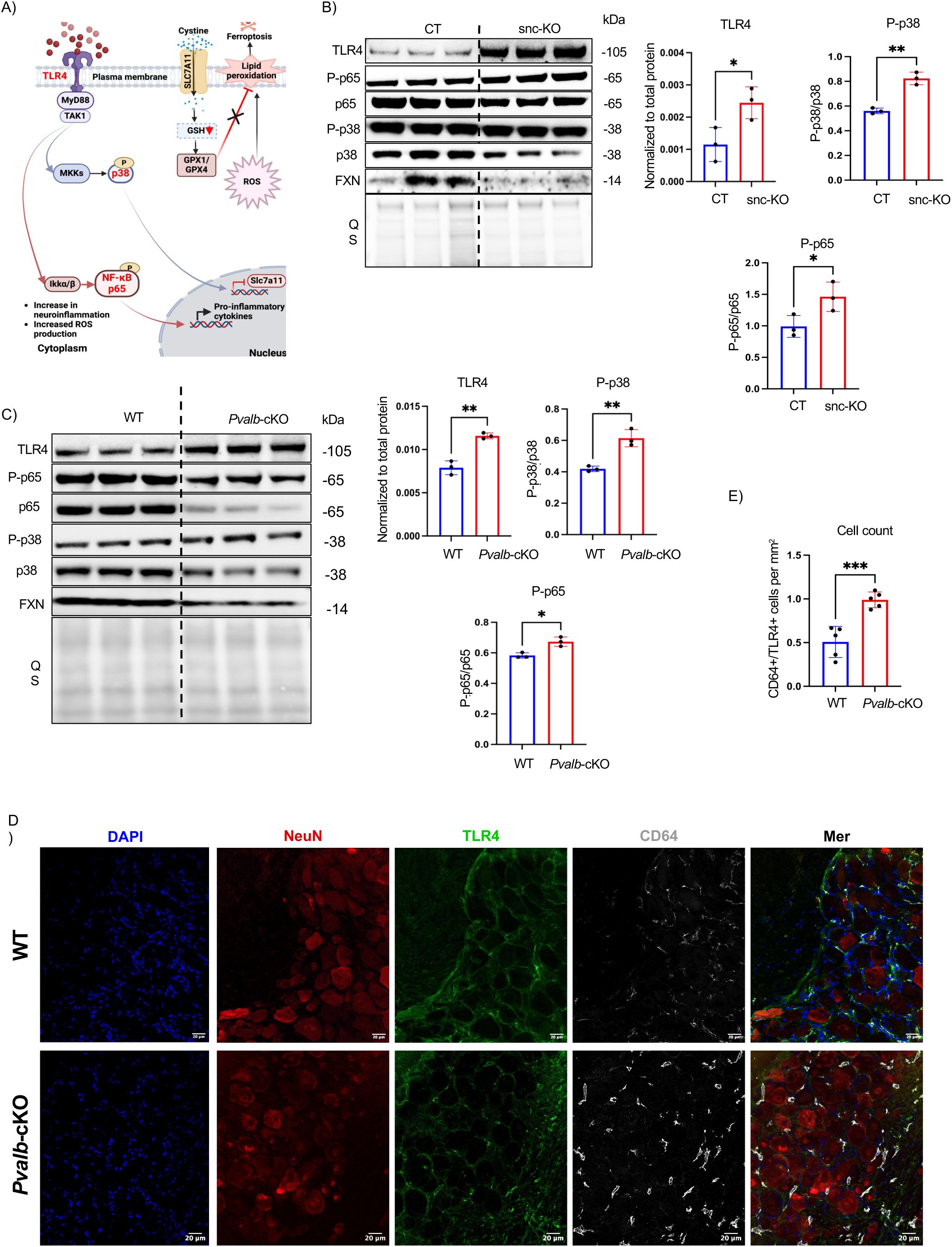
Activation of TLR4 signaling in frataxin-deficient primary sensory neurons culture and dorsal root ganglia. **(A)** Schematic representation of canonical TLR4 signaling pathways and their relationship to oxidative stress and inflammatory responses. TLR4 activation leads to downstream signaling through MyD88/TAK1, resulting in phosphorylation of p38 MAPK and NF-κB p65, induction of pro-inflammatory cytokine production, and increased reactive oxygen species (ROS) production. Interactions with redox pathways involving GPX1/GPX4 and the cystine/glutathione axis are illustrated. **(B)** Immunoblot analysis of TLR4 pathway activation in primary sensory neuron cultures following frataxin depletion (snc-KO) compared to control (CT) conditions. Representative blots and quantification of TLR4, phosphorylated p38 (P-p38), total p38, phosphorylated NF-κB p65 (P-p65), total p65, and frataxin (FXN) are shown. Protein levels were normalized to total protein (QS loading control). Data are presented as mean ± SEM from independent biological replicates (n=3); *P < 0.05; **P < 0.01 (unpaired two-tailed Student’s t-test). **(C)** Immunoblot analysis of TLR4 pathway activation in lumbar (L1–L5) DRG lysates from control and post-symptomatic *Pvalb-*cKO mice. Representative blots and quantification of TLR4, P-p38/p38 ratio, and P-p65/p65 ratio are shown. Protein levels were normalized to total protein. Data represent mean ± SEM; statistical significance determined using unpaired two-tailed Student’s t-test; *P < 0.05; **P < 0.01. **(D)** Immunofluorescence analysis of DRG sections from control and *Pvalb*-cKO mice stained for DAPI (blue), NeuN (neuronal marker, red), TLR4 (green), and CD64 (myeloid marker, white). TLR4 immunoreactivity is predominantly detected in small non-neuronal cells within the DRG microenvironment and shows substantial overlap with CD64-positive cells. Scale bars: 20 μm. **(E)** Quantification of CD64⁺/TLR4⁺ cells in DRG tissue sections, expressed as cells per mm². Data represent mean ± SEM from biological replicates (n=5 in each genotype); ***P < 0.001 (unpaired two-tailed Student’s t-test).

Immunoblot analysis of primary sensory neuron cultures revealed a significant increase in TLR4 protein levels in FXN-deficient neurons compared to controls (Figure 2B). To determine whether the elevated TLR4 expression was accompanied by downstream pathway activation, we quantified phosphorylation of p38 MAPK and NF-κB p65, two established mediators of TLR4 signaling[36, 38–42]. Both phosphorylated p38 and phosphorylated p65 levels were significantly increased in FXN-deficient neurons, consistent with activation of TLR4-associated signaling pathways (Figure 2B).

We next examined whether TLR4 activation could be detected *in vivo* in the *Pvalb*-cKO model. Despite the previously noted dilution effect in whole DRG lysates, immunoblot analysis demonstrated increased TLR4 expression and elevated p38 and p65 phosphorylation at the post-symptomatic stage (Figure 2C). In contrast, no significant activation was detected at the pre-symptomatic stage (Figure S3A), indicating that TLR4 pathway engagement emerges during disease progression.

To determine the cellular source of TLR4 activation within the DRG, we performed immunofluorescence analysis on tissue sections. TLR4 immunoreactivity was readily detectable at the post-symptomatic stage, whereas minimal signal was observed in pre-symptomatic tissue (Figure 2C and Figure S3B). Notably, TLR4 signal did not seem to co-localize with the neuronal markers NeuN. Instead, TLR4 expression was primarily observed in smaller non-neuronal cells within the DRG microenvironment (Figure 2D).

Given that TLR4 is well-characterized innate immune receptor, we next assessed whether TLR4-positive cells corresponded to immune populations. Co-immunostaining with the myeloid marker CD64 demonstrated that TLR4 expression predominantly localized to CD64-positive cells (Figure 2D). Quantitative analysis demonstrated a significant increase in the proportion of CD64⁺/TLR4⁺ cells in post-symptomatic *Pvalb*-cKO DRGs compared to controls (Figure 2E), accompanied by morphological features consistent with immune cell activation (rounded structure with larger soma) (Figure S3B). Flow cytometric analysis further confirmed an increased frequency of CD64⁺/TLR4⁺ cells in *Pvalb*-cKO DRGs at both pre and post-symptomatic stages (Figure S4A,B).

Collectively, these findings indicate that TLR4 pathway activation in frataxin-deficient DRGs is predominantly associated with immune cells within the tissue microenvironment and becomes progressively amplified during disease progression.

### Macrophages exhibit enhanced TLR4-associated inflammatory signatures in frataxin-deficient DRGs

To further characterize immune populations contributing to TLR4 signaling, we interrogated immune cell subsets identified in the scRNA-seq dataset. Clustering analysis resolved 4 major immune populations based on canonical marker expression: neutrophils, macrophages, monocytes, proliferating (Mki67+) cells (Figure 3A, B; S5A).

**Figure 3:**
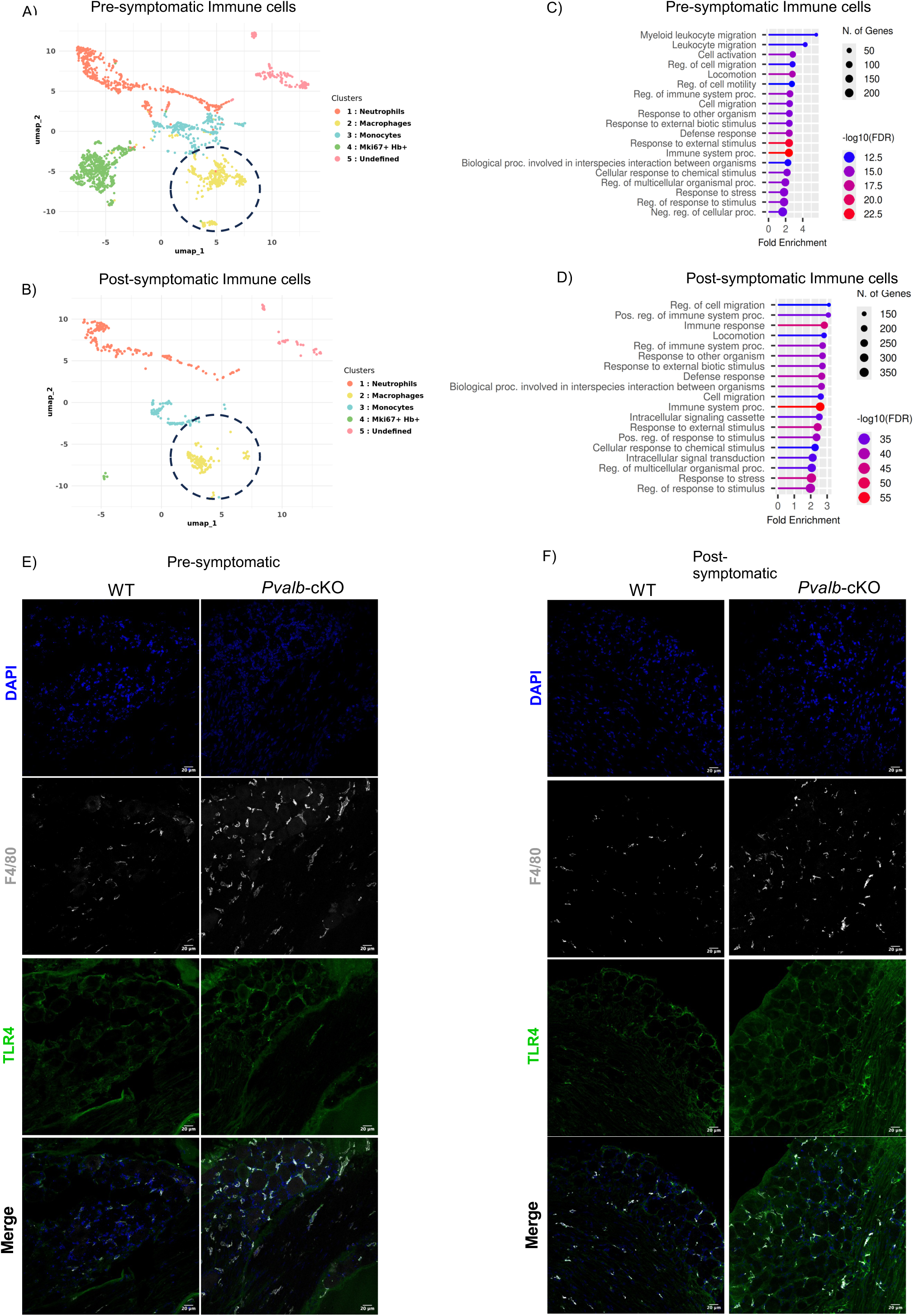
Progressive macrophage expansion and inflammatory activation in *Pvalb*-cKO DRGs. **(A–B)** UMAP representation of immune cell subsets extracted from DRG scRNA-seq datasets at the pre-symptomatic (A) and post-symptomatic (B) stages. Distinct immune populations were identified based on canonical markers, including neutrophils, macrophages, monocytes, proliferating (Mki67⁺) cells, and an undefined cluster. The dashed circle indicates the macrophage cluster selected for downstream analyses. **(C–D)** Gene Ontology (GO) biological process enrichment analysis of differentially expressed genes in the macrophages cluster at the pre-symptomatic (C) and post-symptomatic (D) stages. Enriched terms include leukocyte migration, immune system processes, cellular activation, and inflammatory responses. **(E–F)** Immunofluorescence staining of DRG sections from control and *Pvalb*-cKO mice at pre-symptomatic (E) and post-symptomatic (F) stages. Sections were stained for DAPI (blue), F4/80 (macrophage marker, white), and TLR4 (green). Scale bars: 20 μm.

Given previous evidence implicating macrophages in neuroinflammatory responses associated with Friedreich ataxia ganglionopathy[43], we focused on this population (Figure 3A,B yellow cluster delimitated/highlighted with dash line & Figure S5B). Differential expression analysis identified *Tlr4* among the significantly upregulated genes in macrophages from *Pvalb*-cKO mice compared to controls at both disease stages.

Gene Ontology (GO) enrichment analysis on differentially expressed genes within the macrophage cluster revealed enrichment of pathways associated with stress responses, immune activation, inflammatory signaling, leukocyte migration, and cellular activation at both disease stages. Notably, enrichment magnitude and statistical significance were increased at the post-symptomatic stage, consistent with progressive amplification of inflammatory programs (Figure 3C,D).

To determine whether TLR4 protein expression localized to macrophages *in vivo*, we performed co-immunostaining of TLR4 with the macrophage marker F4/80 in DRG sections. TLR4 immunoreactivity overlapped with F4/80-positive cells at both pre-symptomatic and post-symptomatic stages (Figure 3E,F). Collectively, these findings identify macrophages as a prominent TLR4-expressing immune cell population exhibiting enhanced inflammatory signatures in *Pvalb*-cKO DRGs.

### Satellite glial cell activation and S100a8 upregulation in *Pvalb*-cKO DRGs

Following the identification of macrophage-associated TLR4 activation, we next investigated potential upstream signals within the DRG microenvironment that could contribute to this response. Given the abundance and close anatomical association of SGCs within sensory neurons, we re-analyzed the scRNA-seq dataset to specifically interrogate SGC populations at both disease stages.

UMAP clustering resolved distinct SGC subpopulations at each disease stage (Figure S6 A,B). Feature plots confirmed robust expression of canonical SGC markers, including *Fabp7* and *S100b*, validating cluster identity (Figure S6 C,D). Differential expression analysis followed by GO enrichment revealed significant enrichment of pathways associated with cellular activation, protein localization, metabolic regulation, and stress responses in pre-symptomatic SGCs (Figures 4A; S6 E,F). At the post-symptomatic stage, SGCs exhibited enrichment of translation, macromolecular biosynthesis, intracellular transport, and broader immune- and stress-associated processes (Figure 4B; S6 G,H), consistent with sustained and amplified glial activation.

**Figure 4:**
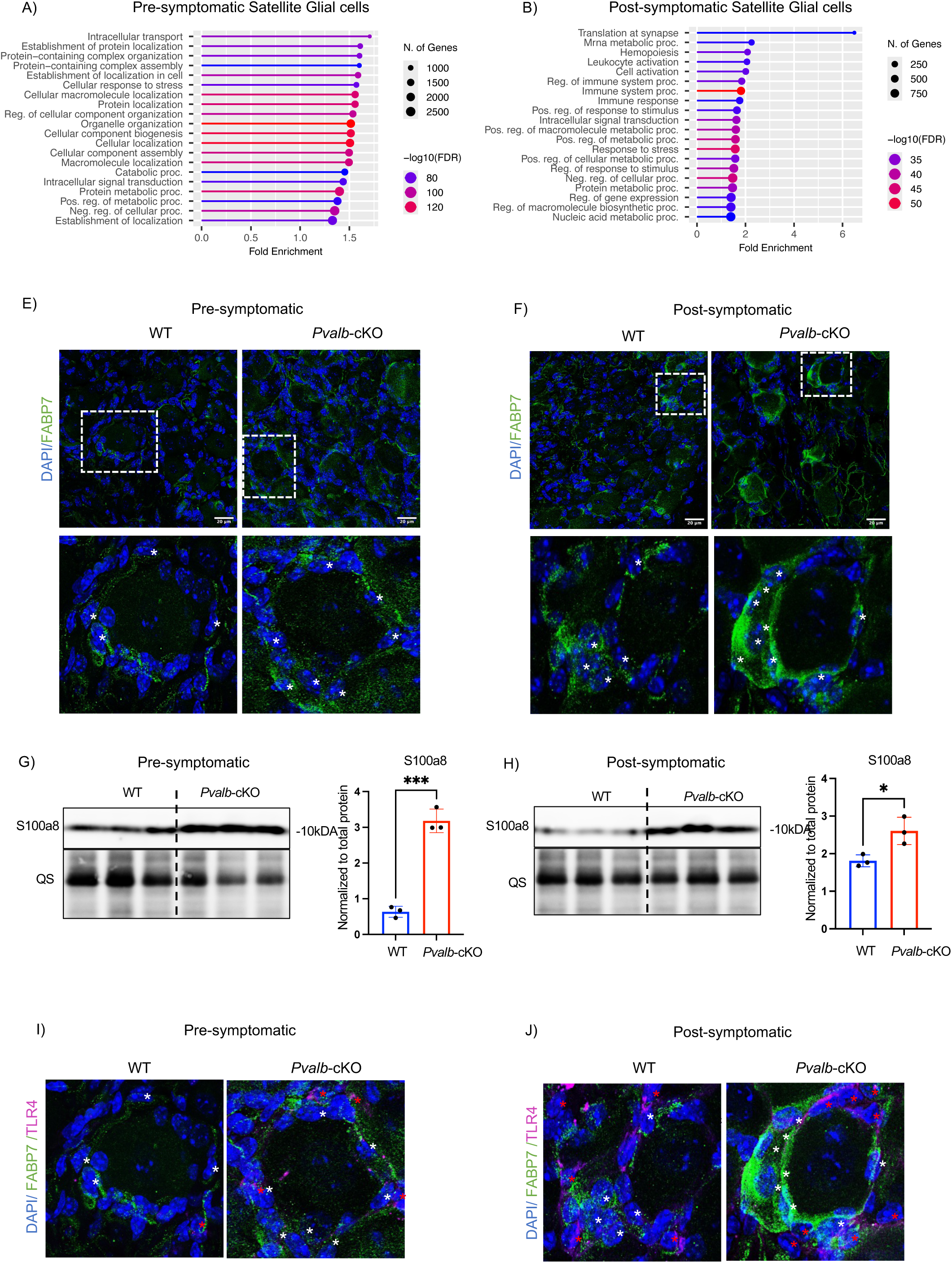
Satellite glial cell activation and S100a8 upregulation in *Pvalb*-cKO DRGs. **(A–B)** GO biological process enrichment analysis of differentially expressed genes in satellite glial cells (SGCs) at pre-symptomatic (A) and post-symptomatic (B) stages. Enriched terms include protein localization and intracellular organization at pre-symptomatic stages, and translation, immune-related processes, and inflammatory signaling at post-symptomatic stages. **(E–F)** Immunofluorescence staining of DRG sections from control and *Pvalb*-cKO mice for FABP7 (SGC marker, green) and DAPI (blue) at pre-symptomatic (E) and post-symptomatic (F) stages. Asterisks indicate SGC surrounding neuronal soma. Insets show higher magnification views. Scale bars: 20 μm & 10 µm (zoomed in section) **(G–H)** Immunoblot analysis of S100a8 expression in DRG lysates from control and *Pvalb*-cKO mice at pre-symptomatic (G) and post-symptomatic (H) stages. Protein levels were normalized to total protein (QS loading control). Data are presented as mean ± SEM from biological replicates (n=3); *P< 0.05, ***P < 0.001 (unpaired two-tailed Student’s t-test) **(I–J)** Co-immunostaining for FABP7 (green) and TLR4 (magenta). Asterisks indicate FABP7-positive satellite glial cells (white) and TLR4-positive cells (red), respectively. Scale bars: 10µm

To validate these transcriptional changes *in vivo*, we performed immunofluorescence analysis using FABP7 as a marker of activated SGCs. *Pvalb*-cKO DRGs displayed increased FABP7 immunoreactivity at both pre- and post-symptomatic stages (Figure 4E,F).

Among the genes differentially expressed in SGCs, *S100a8* was consistently upregulated at both disease stages (Figure S6 I,J). *S100a8* encodes a damage-associated molecular pattern (DAMP) protein capable of engaging TLR4 signaling. Its increased expression in SGCs (Figure 4G,H) is therefore consistent with a potential role in modulating TLR4-assocated immune response within the DRG microenvironment (Figure 4I,J).

Collectively, these findings indicate that SGCs in *Pvalb*-cKO DRG undergo progressive transcriptional activation accompanied by upregulation of S100a8 expression, supporting a non–cell-autonomous model in which glial activation may contribute to macrophage-associated TLR4 signaling during disease progression.

### Pharmacological inhibition of TLR4 ameliorates soma size and mitochondrial deficits in FXN-deficient sensory neurons

Given the consistent activation of TLR4 receptor observed in both frataxin deficient primary sensory neurons and *Pvalb*-cKO DRGs, we next examined whether this pathway functionally contributes to the neuronal deficits associated with frataxin loss. To address this, we employed a pharmacological approach to inhibit TLR4 signaling.

TAK-242 (Resatorvid) is a selective small-molecule inhibitor of TLR4 that binds its intracellular domain and disrupts interaction with adaptor proteins including MyD88 and TRIF[44–48], thereby blocking downstream NF-κB and MAPK signaling pathways [49].

Consistent with our previous characterization of primary DRG cultures, FXN-deficient neurons exhibited reduced soma size and axonal thinning compared to controls[50]. Treatment with TAK-242 significantly increased soma size and improved axonal morphology relative to untreated FXN-deficient neurons (Figure 5A). These findings indicate that TLR4 signaling contributes to the structural alterations observed in frataxin-deficient sensory neurons.

**Figure 5:**
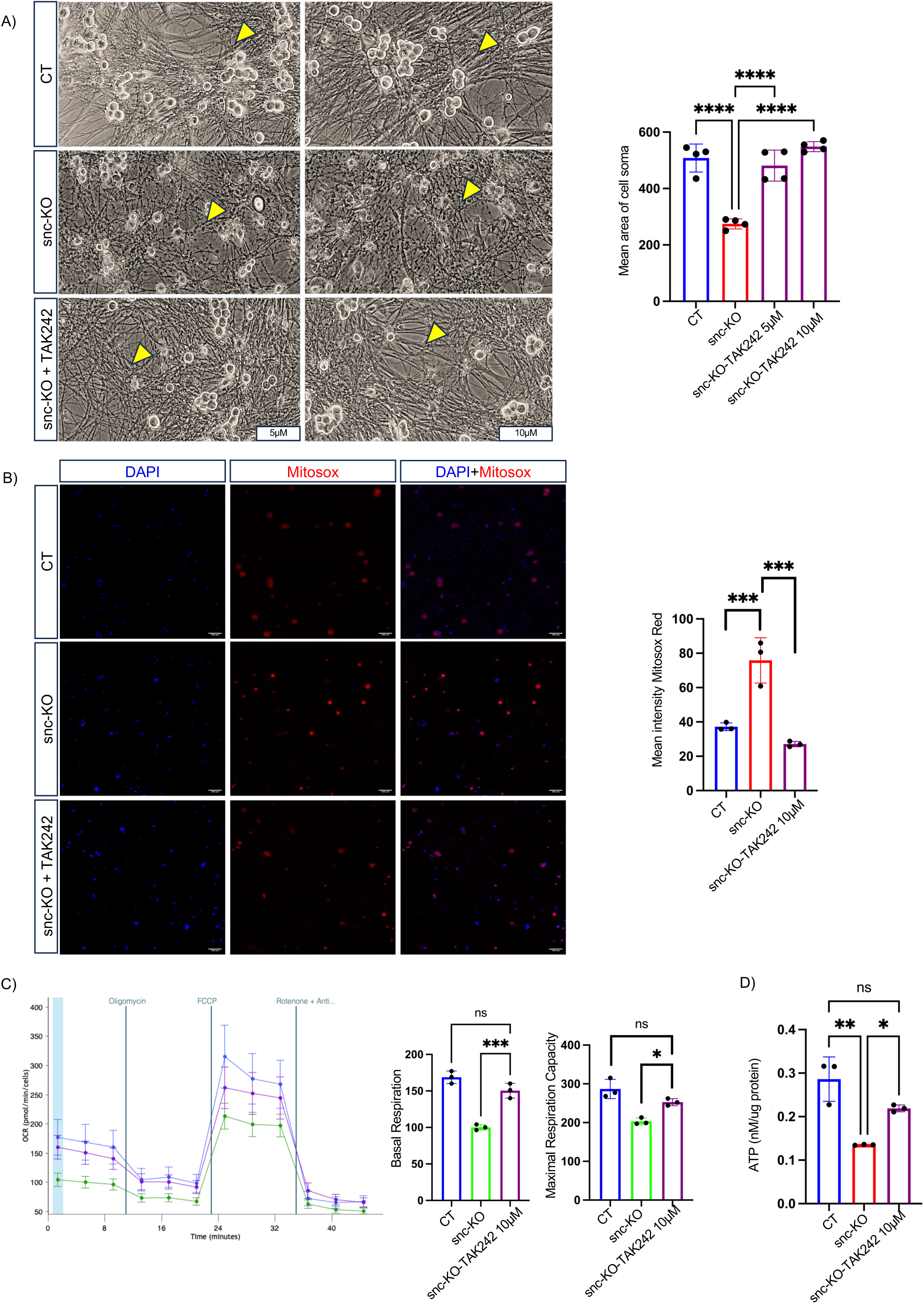
Pharmacological inhibition of TLR4 with TAK-242 in FXN-deficient primary sensory neurons. **(A)** Representative brightfield images of primary sensory neuron cultures from control (CT) and FXN-deficient neurons (snc-KO) treated with vehicle or TAK-242 (5 µM and 10 µM). Yellow arrowheads indicate neuronal axons soma. Quantification of mean neuronal soma area is shown on the right. Data represent mean ± SEM from independent cultures (n=3). **(B)** Representative fluorescence images of mitochondrial superoxide production using MitoSOX Red staining. Nuclei were counterstained with DAPI (blue). Quantification of mean MitoSOX Red intensity is shown on the right. Data represent mean ± SEM from independent experiments (n=3). (**C)** Mitochondrial respiration analysis in primary sensory neuron cultures measured using a Seahorse extracellular flux assay. Oxygen consumption rate (OCR) was monitored following sequential injections of oligomycin, FCCP, and rotenone/antimycin A. Quantification of basal respiration and maximal respiration capacity is shown on the right. Data represent mean ± SEM from independent experiments (n=3). **(D)** Quantification of cellular ATP levels in primary sensory neuron cultures. Data are presented as mean ± SEM from independent biological replicates n=3). For all quantifications, statistical significance was determined using one-way analysis of variance (ANOVA) followed by post hoc multiple comparison testing. Significance levels are indicated as: \**P* < 0.05*, **P < 0.01, ***P < 0.001*, and \*\*\*\**P* < 0.0001; ns, not significant. Scale bars: 100 µm.

Given the link between frataxin deficiency, TLR4 activation, and oxidative stress pathways, we next assessed mitochondrial reactive oxygen species (ROS) using MitoSOX probe. FXN-deficient neurons showed elevated mitochondrial ROS levels compared to controls (Figure 5B). TAK-242 treatment significantly reduced ROS accumulation, consistent with attenuation of oxidative stress.

To determine whether these changes translated into improved mitochondrial function, we performed Seahorse metabolic flux analysis. FXN-deficient neurons exhibited reduced basal and maximal respiratory capacity compared with controls (Figure 5C). Pharmacological inhibition of TLR4 partially restored both parameters. In parallel, intracellular ATP levels were significantly increased in TAK-242-treated FXN-deficient neurons compared to untreated counterparts (Figure 5D).

Collectively, these data indicate that TLR4 signaling contributes to structural and metabolic impairments in frataxin-deficient sensory neurons and that its pharmacological inhibition mitigates these deficits *in vitro*.

### TLR4 inhibition with TAK-242 delays proprioceptive neuron degeneration in *Pvalb*-cKO mice

To determine whether TLR4 signaling contributes to disease progression *in vivo*, we evaluated the effects of pharmacological TLR4 inhibition using TAK-242 in *Pvalb-*cKO mice. TAK-242 (10 mg/kg) was administered intraperitoneally starting at the pre-symptomatic stage (3.5 weeks), and behavioral and electrophysiological assessments were monitored as the animals progressed toward the post-symptomatic stage (7.5–10.5 weeks) (Fig. 6A).

**Figure 6:**
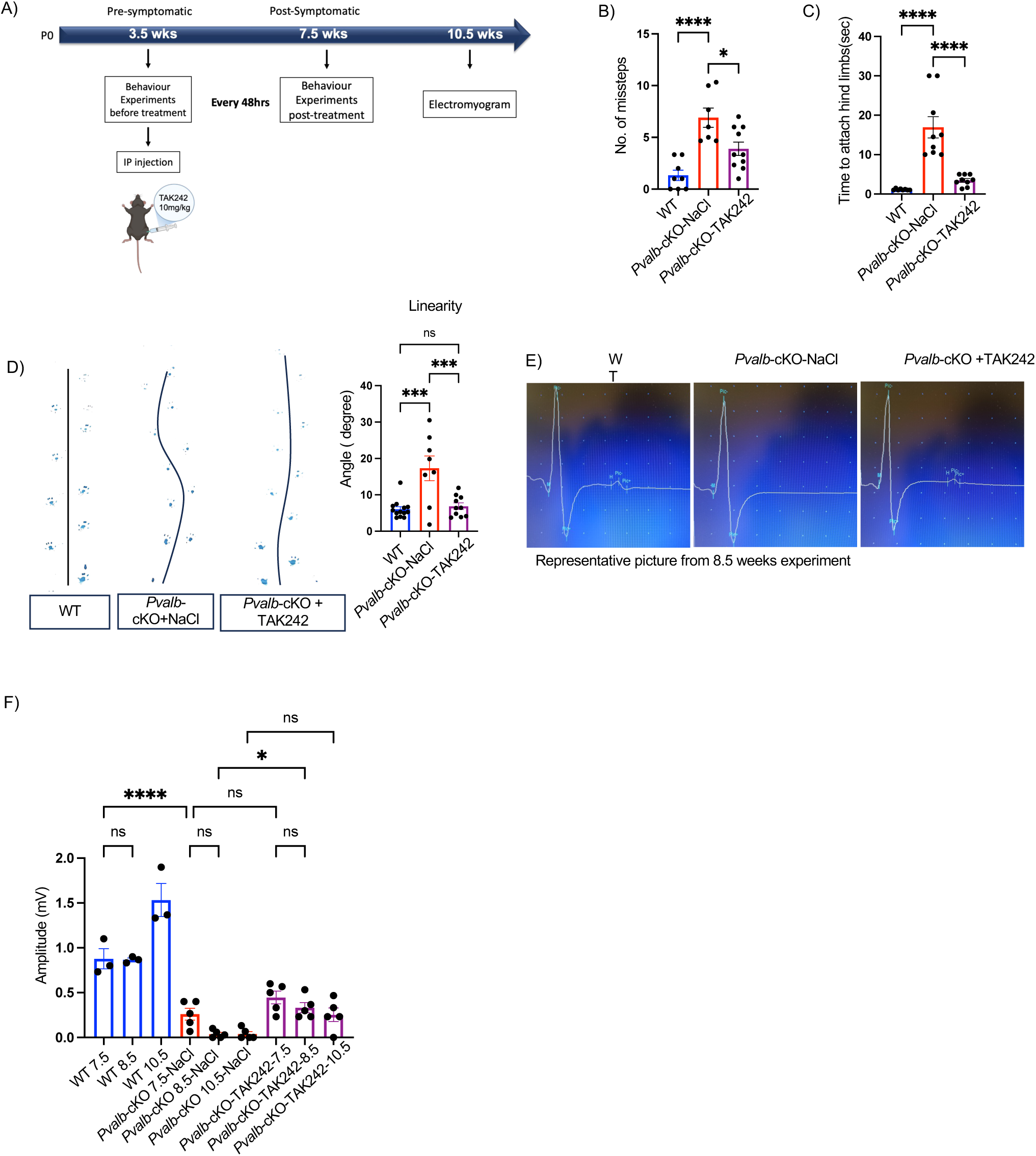
Pharmacological inhibition of TLR4 with TAK-242 mitigates motor and electrophysiological deficits in FXN-Pvalb-cKO mice. **(A)** Experimental design of the TAK-242 treatment paradigm. FXN-Pvalb-cKO mice received intraperitoneal (IP) injections of the TLR4 inhibitor TAK-242 (10 mg/kg) beginning at the pre-symptomatic stage (3.5 weeks of age) and subsequently every 48 hours. Behavioural assessments were performed prior to treatment and at the post-symptomatic stage of 7.5 weeks. Electromyography (EMG) recordings were performed at 7.5, 8.5 and 10.5 weeks of age. **(B)** Quantification of missteps during motor coordination test on crenelated bar test. **(C)** Time required to attach hind limbs (sec) on wire hanging test. **(D)** Gait analysis and linearity measurements. Representative trajectory traces of hind-limb movement in WT, FXN-*Pvalb*-cKO, and TAK-242–treated *Pvalb*-cKO mice are shown. Quantification of the hind-limb angle deviation (linearity) is shown. **(E)** Representative EMG recordings obtained at 8.5 weeks from WT, *Pvalb*-cKO-NaCl saline-treated, and *Pvalb*-cKO mice treated with TAK-242. **(F)** Quantification of EMG signal amplitude (mV) across disease stages (7.5, 8.5, and 10.5 weeks). Data are presented as mean ± SEM; For behavioural test: n=8 for WT, n=7 for *Pvalb*-cKO-NaCl, n=10 for TAK-242–treated *Pvalb*-cKO.For EMG: n=3 for WT, n=5 for *Pvalb*-cKO-NaCl, n=5 for TAK-242–treated *Pvalb*-cKO. Statistical analysis was performed using one-way ANOVA followed by post hoc multiple comparisons. Significance levels are indicated as: \**P* < 0.05*, **P < 0.01, ***P < 0.001*, and \*\*\*\**P* < 0.0001; ns, not significant.

As previously reported [16], *Pvalb*-cKO mice develop progressive motor and sensorimotor deficits. Consistent with this established phenotype, untreated *Pvalb*-cKO mice exhibited increased number of missteps on the Notched bar test (Fig. 6B) and prolonged latency in the hanging wire test (Fig. 6C). TAK-242 treatment significantly reduced the number of missteps and improved performance in the hanging wire assay compared to untreated *Pvalb*-cKO mice (Fig. 6B-C, Supplementary video 1). Similarly, gait abnormalities previously described in this model were recapitulated, with *Pvalb*-cKO mice displaying increased in hind-limb angle deviation (linearity) relative to WT control (Fig. 6D). Treatment with TAK-242 improved gait parameters and reduced angular deviation toward WT levels. Electrophysiological analysis further confirmed the previously reported progressive decline of the sensorimotor reflex (H-wave) in *Pvalb*-cKO mice, with near-complete loss by approximately 8.5 weeks of age (Fig. 6E-F). In contrast, TAK-242-treated mice showed partial preservation of H-wave amplitude, which remained detectable up to 10.5 weeks of age (Fig. 6E-F).

Collectively, these findings indicate that pharmacological inhibition of TLR4 signaling delays the progressive motor and sensorimotor deficits in *Pvalb*-cKO mice, supporting a functional contribution of TLR4-mediated inflammatory signaling to disease progression *in vivo*.

## Discussion

Friedreich ataxia (FA) is characterized by progressive degeneration of large proprioceptive sensory neurons (pSNs) within the dorsal root ganglia (DRG), leading to severe sensory ataxia. Although frataxin deficiency is well established to impair mitochondrial function and redox homeostasis, the mechanisms linking neuron-intrinsic metabolic dysfunction to tissue-level degeneration remain incompletely understood. Here, we identify TLR4-associated neuroimmune signaling within the DRG microenvironment as a contributor to disease progression in a neuron-specific frataxin-deficient mouse model (*Pvalb*-cKO).

Single-cell RNA sequencing revealed transcriptional alterations in frataxin-deficient pSNs consistent with mitochondrial dysfunction and sustained cellular stress. In particular, downregulation of genes involved in redox regulation and mitochondrial homeostasis, including *Gpx1* and *Ucp2*, together with upregulation of stress-responsive genes such as *Hsp90aa1*, *Hsp90ab1*, and *Camkk2*, is consistent with a persistent stress state. These findings align with the established role of frataxin in mitochondrial function and support the intrinsic metabolic vulnerability of pSNs in FA.

A central finding of this study is that neuron-restricted frataxin deficiency is accompanied by substantial transcriptional and functional changes in non-neuronal cell populations within the DRG. Despite preserved frataxin expression, both macrophages and satellite glial cells (SGCs) exhibited robust activation signatures, highlighting the importance of non-cell-autonomous mechanisms in disease progression. Macrophages displayed enrichment of inflammatory pathways and increased expression of TLR4, together with an increased abundance of CD64⁺/TLR4⁺ immune cells *in vivo*, indicating activation of the innate immune compartment.

SGCs also exhibited pronounced transcriptional activation, with enrichment of pathways related to metabolic regulation, protein localization, and stress responses. Notably, *S100a8*, encoding a damage-associated molecular pattern (DAMPs) capable of activating TLR4 signaling, was significantly upregulated in SGCs at both disease stages. These observations support a model in which activated SGCs contribute to the inflammatory microenvironment of the DRG and may modulate immune cell activation through the release of endogenous TLR4 ligands.

Neuroinflammatory processes in FA have been suggested in previous studies, primarily in central nervous system regions. Evidence from patient-derived samples, imaging studies, and experimental models indicates the presence of inflammatory signatures and glial activation, including microglial activation in the cerebellum and brainstem, and systemic immune alterations detectable in peripheral tissue [51–54]. Importantly, neuropathological analyses of patient DRGs have reported SGCs abnormalities and inflammatory features associated with neuronal degeneration, supporting the relevance of neuroimmune interactions in the peripheral nervous system [7]. TLR4-associated signaling has also been reported in patient-derived fibroblasts, although these observations remain limited [55]. However, the cellular and molecular mechanisms underlying these responses have remained poorly defined.

An important aspect of the present study is that frataxin deficiency is restricted to proprioceptive neurons, while non-neuronal cells retain normal frataxin levels. This experimental design allows the identification of non-cell-autonomous mechanisms linking neuronal stress to microenvironmental responses. In contrast, in patients and in other FA models, frataxin deficiency affects multiple cell types, suggesting that both cell-intrinsic and non-cell-autonomous mechanisms contribute to disease progression. Our findings therefore define a pathway by which neuron-intrinsic metabolic dysfunction can initiate and propagate inflammatory signaling within the DRG microenvironment.

In line with this observation, TLR4 signaling was activated at both cellular and tissue levels, as evidence by increased TLR4 expression and phosphorylation of downstream signaling mediators p38 MAPK and NF-κB p65. Importantly, *in vivo* TLR4 expression was predominantly localized to CD64-positive immune cells, indicating that TLR4-mediated signaling arises primarily within immune compartment. These findings position TLR4 as a potential integrator linking neuronal stress to immune activation.

The functional relevance of this pathway was supported by pharmacological inhibition experiments. In primary sensory neuron cultures, inhibition of TLR4 signaling improved neuronal morphology, reduced mitochondrial reactive oxygen species, and partially restored mitochondrial function. *In vivo*, administration of TAK-242 from the pre-symptomatic stage delayed the progression of motor and sensorimotor deficits in *Pvalb*-cKO mice, including improvements in motor coordination, gait, and preservation of sensorimotor reflexes. Together, these results indicate that TLR4-associated pathways contribute to disease progression and that their modulation can influence disease trajectory.

TLR4-mediated inflammatory signaling has been implicated in multiple neurodegenerative disorders, including Alzheimer’s disease, Parkinson’s disease and sensory neuropathies [56–60] [43], where it contributes to chronic neuroinflammation and neuronal dysfunction. Our findings extend this paradigm to FA and suggest that neuroimmune interactions represent an additional layer of pathophysiology beyond primary mitochondrial impairment, with TLR4 acting as a critical node linking metabolic stress to inflammatory amplification within the DRG.

Collectively, our data support a model in which frataxin deficiency in proprioceptive neurons initiates metabolic stress that triggers activation of SGCs and immune cells within the DRG microenvironment. This non-cell-autonomous response is associated with increased production of inflammatory mediators and TLR4 ligands, including S100A8, leading to amplification of TLR4-dependent signaling pathways that exacerbate neuronal dysfunction. Pharmacological inhibition of TLR4 mitigates several aspects of this pathology, highlighting this pathway as a potential therapeutic target in FA.

Several limitations should be considered. Although our data identify TLR4-assocaited signaling as a contributor to disease progression, the precise molecular mechanisms linking neuronal stress, SGC activation, and macrophage responses remain to be fully defined. In particular, the cellular sources and regulation of TLR4 ligands within the DRG microenvironment require further investigation. In addition, longer-term and post-symptomatic studies will be necessary to determine whether sustained TLR4 inhibition provides durable neuroprotection and whether similar mechanisms operate in human disease.

In summary, this study identifies TLR4-mediated neuroimmune signaling as a previously unrecognized component of DRG pathology in Friedreich ataxia and provides a conceptual model linking neuron-intrinsic metabolic dysfunction to microenvironment-driven disease progression, highlighting neuroimmune pathways as potential therapeutic targets.

## Materials and Methods

### Animals and tissue collection

All animal procedures were performed in accordance with experimental protocols. Conditional frataxin knockout mice (*Fxn*^L3/L-^; *Pvalb*^tm1(Cre)Arbr/J^), in which frataxin deletion is restricted to parvalbumin-expressing neurons, were used as previsouly described [16]. Lumbar dorsal root ganglia (DRGs; L1-L5) were collected at pre-symptomatic (3.5 weeks) and post-symptomatic (7.5 weeks) stages. For each time point, six mice were used (n=3 per genotype).

Mice were anaesthetized with isoflurane and euthanized by cervical dislocation. The lumbar vertebral column was isolated, and a rostral-caudal bisection was performed to expose the spinal cord. The spinal cord and associated axonal projections were removed, and DRGs (L1–L5) were carefully dissected using fine forceps. Surrounding connective tissue and axons were trimmed prior to downstream processing.

### TAK-242 treatment

For in vivo pharmacological studies, TAK-242 (Resatorvid) was administered intraperitoneally at 10 mg/kg every 48 hours for 4-6 weeks, starting at 3.5 weeks of age. TAK242 was diluted in saline containing 0.5% DMSO and untreated mice were injected with saline containing 0,5% DMSO. Sample size: *Pvalb*-cKO TAK242 treated mice (n=10), *Pvalb*-cKO untreated mice (n=7) and untreated control mice (n=8).

### Dissociation of Dorsal Root Ganglia (DRGs)

Freshly isolated DRGs were immediately placed on ice in Hank’s Balanced Salt Solution (HBSS). Tissues were finely minced and subjected to enzymatic digestion at 37°C in a solution containing 20 U/mL of activated papain (PAPL, cat. no. LS003118, Worthington), final concentration of 8mg/mL of Collagenase/Dispase (cat. no. LS004106, Worthington, stock), and 500 µL of TrypLE for 1mL of dissociation buffer (cat. no. 12605036, Life Technologies) for 30–60 minutes in a humidified incubator with 5% CO₂. During the enzymatic digestion, mechanical dissociation was performed by gently triturating the tissue using fire-polished Pasteur pipettes of decreasing diameter. The resulting cell suspension was filtered through a 40 µm cell strainer to remove debris. Cells were further purified using a 5% BSA gradient centrifugation to eliminate dead cells and debris.

### Single-Cell RNA Sequencing

Single-cell RNA sequencing (scRNA-seq) was performed using the 10x Genomics platform at the Cancer Research Center of Lyon (CRCL). Deep sequencing was performed with an average of 100,000 reads per cell at both 3.5 and 7.5-weeks of age time points. All libraries were prepared according to the manufacturer’s protocol using the 10x Genomics Single Cell 3’ RNA Library Preparation Kit. A total of 8,000–10,000 cells per sample were partitioned into nanoliter-scale Gel Bead-in-Emulsions (GEMs) using the 10x GemCode technology. Library construction was completed using 10x Genomics Single Cell 3’ Reagent Kits V3, and sequencing was performed on an Illumina HiSeq 4000 platform, generating 150-bp paired-end reads for each library.

### Sc-RNAseq Data Analysis

Transcriptomic analysis of sensory neurons was performed using single-cell RNA sequencing (scRNA-seq) data processed in Seurat (R, Version 5). Initial pre-processing, including alignment and quantification of reads, was conducted using Cell Ranger. Quality control metrics were applied to exclude low-quality cells, with thresholds set for gene counts, mitochondrial gene content, and unique molecular identifiers (UMIs). Cells passing quality control were normalized using Seurat’s log-normalization method, and the data were scaled to minimize unwanted variability and highly variable genes were identified using the vst method, selecting the top 6,000 features.

Dimensionality reduction was achieved via Principal Component Analysis (PCA), followed by clustering using the Louvain algorithm at multiple resolutions to identify distinct neuronal populations within the dorsal root ganglia (DRG). Proprioceptive sensory neurons (pSNs) were identified based on specific marker genes (*Pvalb, Spp1, Cntnap2, Runx3, Ntrk3*). Differential gene expression analysis between frataxin-deficient and control groups was conducted using the Wilcoxon rank-sum test, with a false discovery rate (FDR) correction and p-values were adjusted for multiple testing using the Benjamini–Hochberg false discovery rate (FDR) correction.

Pathway enrichment analysis was performed using Gene Set Enrichment Analysis (GSEA) and over-representation analysis, implemented with tools such as *fgsea* and *enrichGO*, using gene sets derived from the Gene Ontology (GO) database to identify dysregulated molecular pathways. Visualization was accomplished using UMAP plots, and similar methods were applied to analyze satellite glial cells (SGCs) for non-autonomous effects. All statistical analyses were performed using R scripts.

### Primary sensory neuron culture

Primary DRG sensory neuron culture were cultured as previously published[30].Neurons were derived from mice carrying the conditional frataxin allele and frataxin deletion was induced *in vitro*. All treatments were performed 7 days after frataxin deletion and experiments were performed on DIV28. TAK-242 was used at a final concentration of 10µM based on prior dose optimization experiment.

### Immunofluorescence

DRGs were fixed in 4% paraformaldehyde (PFA) for 1 hour and cryoprotected prior to embedding in Tissue-Tek Cryomatrix. Cryosections (10 µm) were prepared and mounted on slides. Cultured sensory neurons were fixed in 2% PFA for 15 minutes. Samples were permeabilized with 0.2% Triton X-100 in PBS for 20 minutes, washed in PBS, and incubated overnight at 4°C with primary antibodies (Supplementary Table 1) diluted in PBS +0.2% tween (PBST). After washing, secondary antibodies (1:1000) with DAPI (1:1000) were applied at room temperature for 30 minutes. Samples were mounted using Aqua-Poly/Mount (Polysciences 18606). Images were acquired using a Leica TCS SP5X supercontinuum confocal 40X magnification and analyzed with Fiji-ImageJ software.

### Immunoblotting

Cells or DRG tissues were lysed in RIPA buffer and incubated on ice for 20 minutes. Lysates were centrifuged at 11,000 x g for 10 minutes at 4°C, and protein concentration was determined using a Bradford assay. Proteins (5 µg) were labeled using the QuickStain Protein Labeling Kit (Amersham RPN4000), separated by SDS-PAGE, and transferred to nitrocellulose membranes. Total protein signal was used as loading control. Membranes were blocked in 5% BSA in TBST and incubated with primary antibodies (Supplementary Table 2) followed by HRP-conjugated secondary antibodies (1:10,000 to 1:20,000). Signal detection was performed using SuperSignal™ West Femto Maximum Sensitivity Substrate (ThermoFisher 34096) and visualized with the Amersham Imager 600.

### Mitochondrial and morphological analyses

Sensory neurons were seeded in 96-well plates (Greiner 655090) and stained with Hoechst 33342 (1 mg/mL; Merck 94403), MitoSOX (2.5 µM; ThermoFisher M36008) in neurobasal culture medium for 3 hours at 37°C in 5% CO₂. After staining, cells were washed twice in HBSS and fixed in 2% PFA for 15 minutes. Cell survival was quantified by Hoechst segmentation, and soma area was measured using ImageJ. Fluorescence intensity and soma size for each neuron were independently measured through cell segmentation.

### Mitochondrial respiration assay (Seahorse)

Mitochondrial respiration was measured using a Seahorse XF extracellular flux analyzer (Agilent Technologies). Primary sensory neurons were plated in Seahorse XF96 microplates coated with poly-L-lysine, laminin, and Matrigel at a density of 20,000 cells per well and maintained under standard culture conditions. All measurements were performed at DIV28.

Prior to the assay, cells were washed and incubated in Seahorse XF assay medium consisting of XF base medium supplemented with 10 mM glucose, 1 mM sodium pyruvate, and 2 mM L-glutamine (pH 7.4) for 45–60 minutes at 37°C in a non-CO₂ incubator.

Oxygen consumption rate (OCR) was measured under basal conditions followed by sequential injections of oligomycin (1.5 µM), FCCP (0.75 µM), and rotenone/antimycin A (0.5 µM each) to determine ATP-linked respiration, maximal respiration, and non-mitochondrial respiration. OCR measurements were obtained using repeated cycles of mixing, waiting, and measurement according to the manufacturer’s instructions.

Respiration parameters including basal respiration, maximal respiration, and spare respiratory capacity were calculated using Agilent software. OCR values were normalized to cell number.

### Statistical analysis

Data are presented as mean ± SEM. Statistical analyses were performed using GraphPad Prism. Comparisons between two groups were performed using unpaired two-tailed Student’s t-test. Multiple comparisons were analzed using one-way ANOVA followed by appropriated post-hoc tests. A p-value < 0.05 was considered statistically significant.

## Data Availability

The single-cell RNA sequencing data generated in this study will be deposited in a public repository and made available upon publication. Processed data are available from the corresponding author upon reasonable request.

Source data underlying the figures will be provided upon publication. All other data supporting the findings of this study are available from the corresponding author upon reasonable request.

## Author contributions

D.M.C. and H.P. conceived the study. D.M.C. designed and performed experiments, analyzed data, and wrote the manuscript. P.U.A performed bioinformatic analyses and contributed to data interpretation. H.P. supervised the study and contributed to manuscript writing.

## Acknowledgements

We thank Andrea Del Bondi, Beryl Laplace-Builhe, Jessy Van Asperen, Maite Carré Pierrat, Marie Paschaki, Laurence Reutenauer and Cyril Degletagne for their technical assistance and valuable support throughout this study.

This work was supported by BMIC (to D.M.C.), the Association Française pour l’Ataxie de Friedreich (AFAF; to H.P. and M.P.), the Fondation pour la Recherche Médicale (FRM; grant number FDT202304016821 to D.M.C.), and the Friedreich’s Ataxia Research Alliance (FARA; to D.M.C.).

