## Supplementary figures for "TLR4-mediated neuroimmune signalling drives proprioceptive neuron degeneration in Friedreich ataxia"

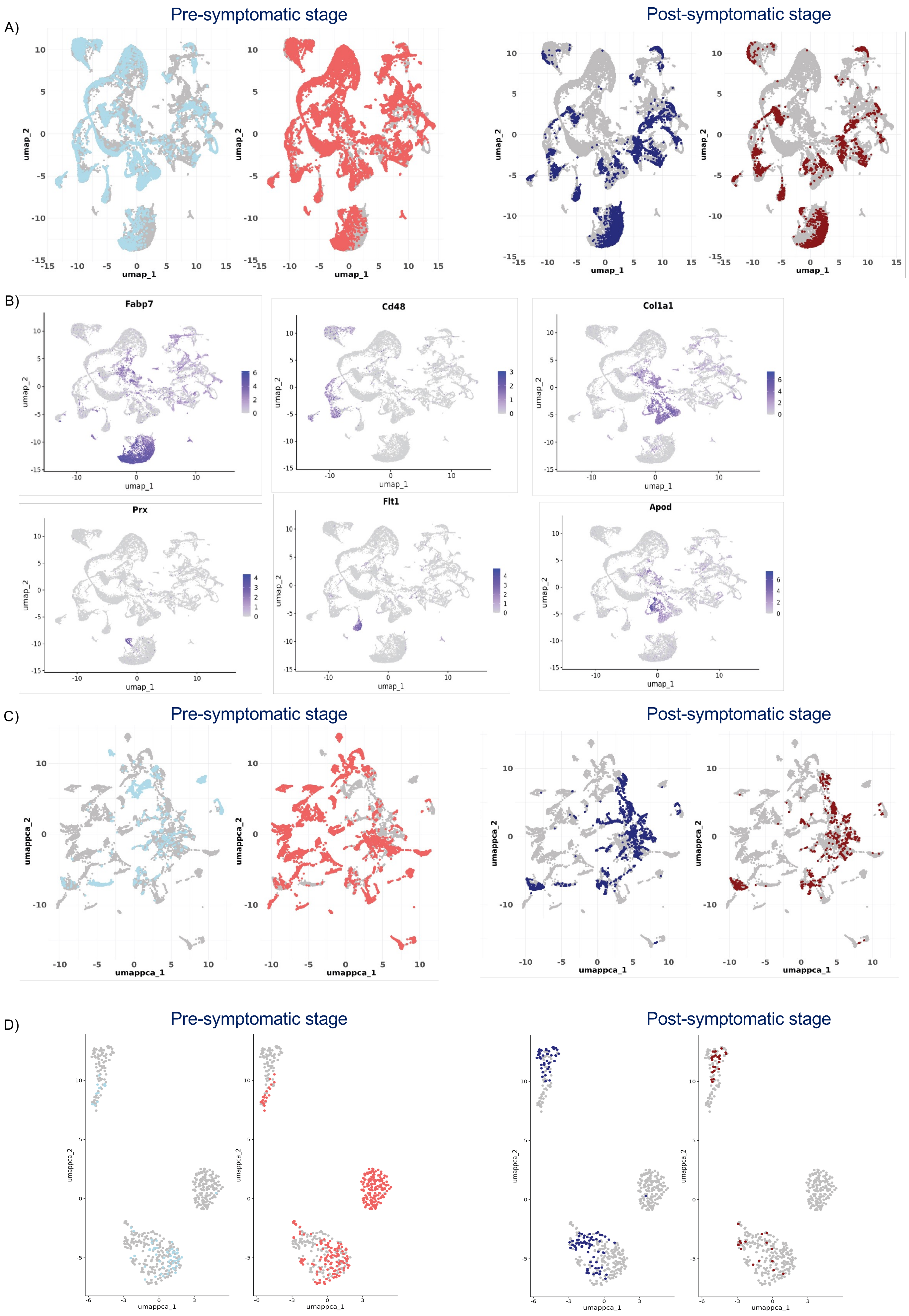
A)
Pre-symptomatic stage
Post-symptomatic stage
umap_1
umap_2
B)
Fabp7
Cd48
Col1a1
Prx
Flt1
Apod
C)
Pre-symptomatic stage
Post-symptomatic stage
umappca_1
umappca_2
D)
Pre-symptomatic stage
Post-symptomatic stage
umappca_1
umappca_2

**Figure S1**

#### Supplementary Figures

##### Figure S1: UMAP visualization of cell populations and marker gene expression across pre-symptomatic and post-symptomatic stages.

**(A)** UMAP embeddings of single-cell transcriptomic data showing the distribution of cells across conditions. Cells corresponding to the pre-symptomatic stage are highlighted in light blue are WT and red (left panels) are *Pvalb-cKO*, while cells from the post-symptomatic stage are highlighted in dark blue are WT and dark red (right panels) are *Pvalb-cKO*. Gray points represent all other cells. **(B)** Feature plots showing expression of selected marker genes across the integrated UMAP embedding, including *Fabp7, Cd48, Col1a1, Prx, Flt1*, and *Apod*. Color intensity (purple gradient) indicates relative gene expression levels. **(C)** UMAP projections highlighting the spatial distribution of neuronal subset assigned to the pre-symptomatic and post-symptomatic stages within the overall data set. Colored cells correspond to the indicated stage, while gray points represent all other cells. **(D)** Subset UMAP plots showing selected pSNs. Cells from pre-symptomatic stage (left panels) and post-symptomatic stage (right panels) are highlighted.

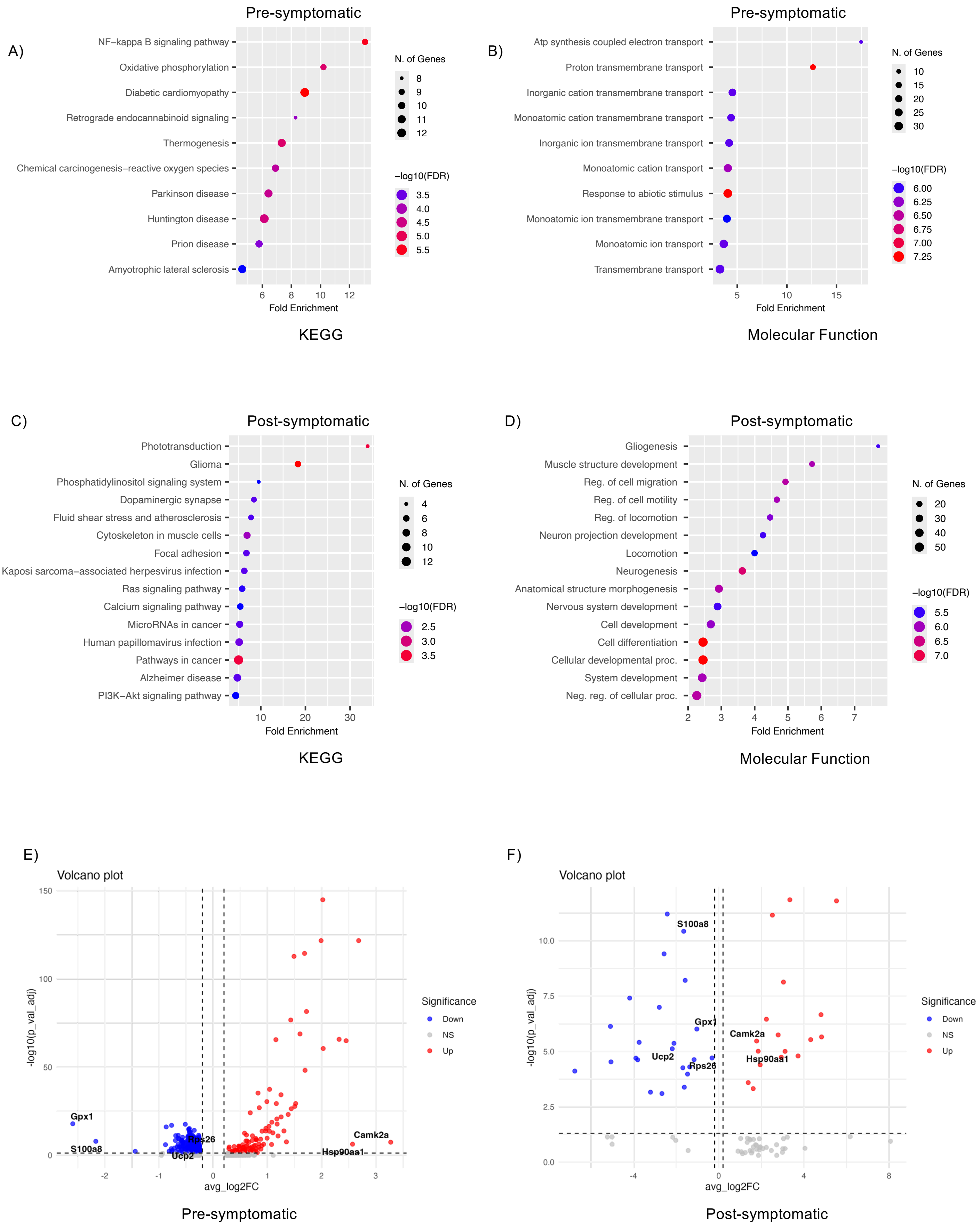
A)
Pre-symptomatic
NF−kappa B signaling pathway
Oxidative phosphorylation
Diabetic cardiomyopathy
Retrograde endocannabinoid signaling
Thermogenesis
Chemical carcinogenesis−reactive oxygen species
Parkinson disease
Huntington disease
Prion disease
Amyotrophic lateral sclerosis
Fold Enrichment
N. of Genes
−log10(FDR)
KEGG
B)
Pre-symptomatic
Atp synthesis coupled electron transport
Proton transmembrane transport
Inorganic cation transmembrane transport
Monoatomic cation transmembrane transport
Inorganic ion transmembrane transport
Monoatomic cation transport
Response to abiotic stimulus
Monoatomic ion transmembrane transport
Monoatomic ion transport
Transmembrane transport
Fold Enrichment
N. of Genes
−log10(FDR)
Molecular Function
C)
Post-symptomatic
Phototransduction
Glioma
Phosphatidylinositol signaling system
Dopaminergic synapse
Fluid shear stress and atherosclerosis
Cytoskeleton in muscle cells
Focal adhesion
Kaposi sarcoma−associated herpesvirus infection
Ras signaling pathway
Calcium signaling pathway
MicroRNAs in cancer
Human papillomavirus infection
Pathways in cancer
Alzheimer disease
PI3K−Akt signaling pathway
Fold Enrichment
N. of Genes
−log10(FDR)
KEGG
D)
Post-symptomatic
Gliogenesis
Muscle structure development
Reg. of cell migration
Reg. of cell motility
Reg. of locomotion
Neuron projection development
Locomotion
Neurogenesis
Anatomical structure morphogenesis
Nervous system development
Cell development
Cell differentiation
Cellular developmental proc.
System development
Neg. reg. of cellular proc.
Fold Enrichment
N. of Genes
−log10(FDR)
Molecular Function
E)
Volcano plot
-log10(p_val_adj)
avg_log2FC
Gpx1
S100a8
Rps26
Ucp2
Camk2a
Hsp90aa1
Significance
Down
NS
Up
Pre-symptomatic
F)
Volcano plot
-log10(p_val_adj)
avg_log2FC
S100a8
Gpx1
Camk2a
Ucp2
Rps26
Hsp90aa1
Significance
Down
NS
Up
Post-symptomatic

**Figure S2**

**Figure S2: Pathway enrichment and differential gene expression analysis in pre-symptomatic and post-symptomatic stages.**

**(A)** KEGG pathway enrichment analysis of genes associated with the pre-symptomatic stage. Dot plots show enriched pathways, including oxidative phosphorylation, NF-κB signaling, thermogenesis, and neurodegeneration-related pathways. The x-axis represents fold enrichment, dot size corresponds to the number of genes contributing to each pathway, and color indicates statistical significance expressed as –log10(FDR). **(B)** Gene Ontology (GO) biological process enrichment analysis for the pre-symptomatic stage. Enriched terms include ATP synthesis-coupled electron transport, proton transmembrane transport, and inorganic/monoatomic ion transmembrane transport. Dot size indicates the number of genes associated with each term, and color represents –log10(FDR). **(C)** KEGG pathway enrichment analysis of genes associated with the post-symptomatic stage. Enriched pathways include phototransduction, phosphatidylinositol signaling, dopaminergic synapse, focal adhesion, and PI3K–Akt signaling. Dot size represents the number of genes contributing to each pathway, while color denotes enrichment significance (–log10(FDR)). **(D)** GO biological process enrichment analysis for the post-symptomatic stage. Enriched terms include gliogenesis, neuron projection development, neurogenesis, locomotion, and cellular developmental processes, suggesting increased involvement of neuronal development and structural remodeling pathways. Dot size represents the number of genes contributing to each pathway, while color denotes enrichment significance (–log10(FDR)). **(E)** Volcano plot showing differential gene expression in the pre-symptomatic stage. The x-axis represents average log2 fold change (avg_log2FC) and the y-axis shows –log10(adjusted p-value). Red points indicate upregulated genes, blue points represent downregulated genes, and gray points indicate non-significant genes. Selected genes of interest are labeled. **(F)** Volcano plot of differentially expressed genes in the post-symptomatic stage using the same thresholds as in (E). Selected genes of interest are labeled.

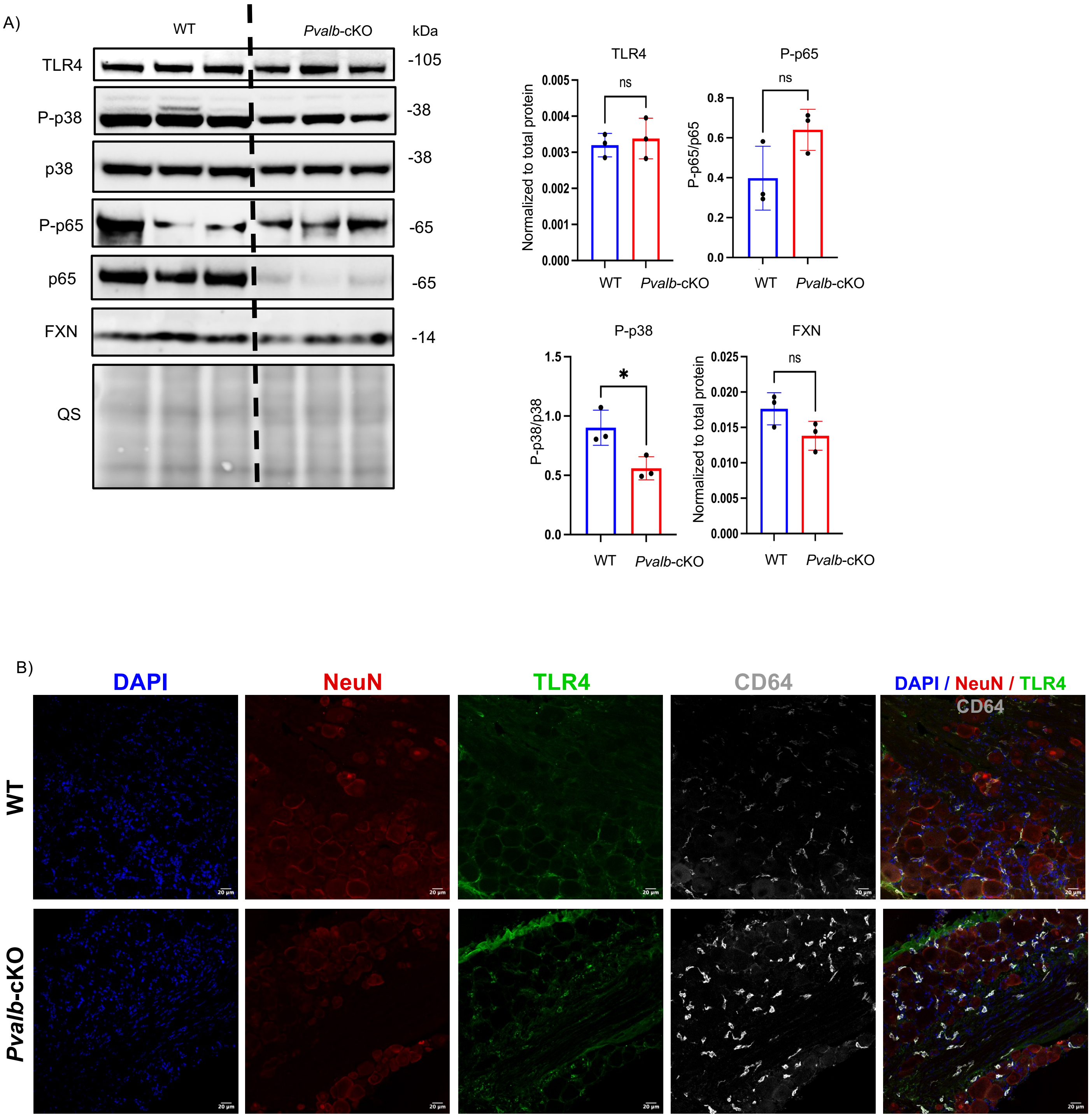
A)
WT
Pvalb-cKO
kDa
TLR4
-105
P-p38
-38
p38
-38
P-p65
-65
p65
-65
FXN
-14
QS
TLR4
ns
Normalized to total protein
0.005
0.004
0.003
0.002
0.001
0.000
WT
Pvalb-cKO
P-p65
ns
P-p65/p65
0.8
0.6
0.4
0.2
0.0
WT
Pvalb-cKO
P-p38
*
P-p38/p38
1.5
1.0
0.5
0.0
WT
Pvalb-cKO
FXN
ns
Normalized to total protein
0.025
0.020
0.015
0.010
0.005
0.000
WT
Pvalb-cKO
B)
DAPI
NeuN
TLR4
CD64
DAPI / NeuN / TLR4
CD64
WT
Pvalb-cKO

**Figure S3**

**Figure S3: TLR4 signaling pathway activation and cellular localization in control and *Pvalb*-cKO mice.**

**(A)** Immunoblot analysis of TLR4 signaling pathway components in control and pre-symptomatic *Pvalb*-cKO samples. Representative blots show protein levels of TLR4, phosphorylated p38 (P-p38), total p38, phosphorylated p65 (P-p65), total p65, and frataxin (FXN). Total protein staining (QS) is shown as a loading control. Protein levels were normalized to total protein levels and phosphorylation is expressed as the ratio of phosphorylated to total protein. Data are expressed as mean ± SEM (n=3);*P $< 0.05$; ns, not significant (unpaired two-tailed Student's t-test). **(B)** Immunofluorescence staining from tissue sections from control and post-symptomatic *Pvalb*-cKO samples. DAPI (blue), NeuN (neuronal marker, red), TLR4 (green), and CD64 (myeloid marker, white). Merged images are shown. Scale bars: 20 μm.

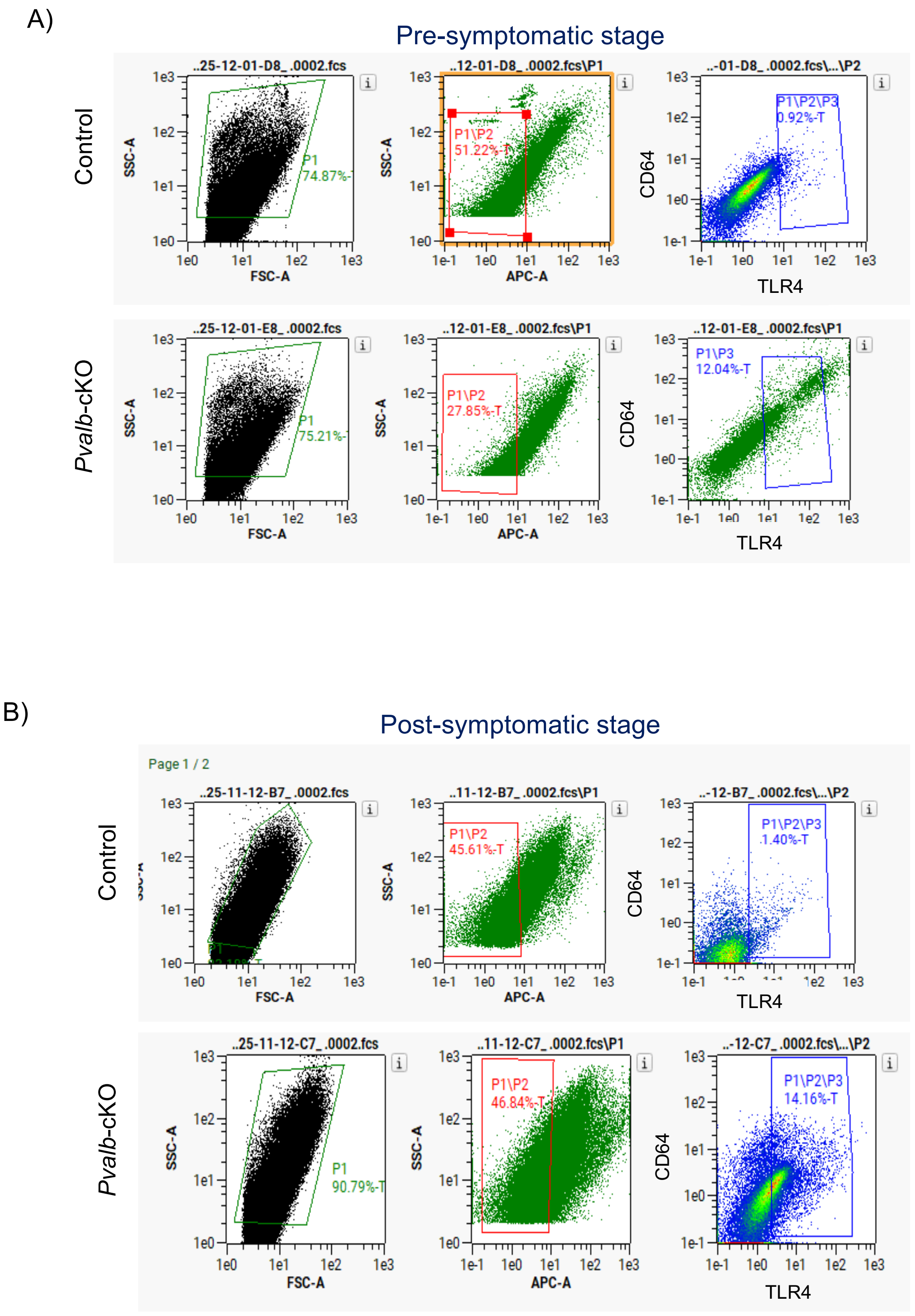
A)
Pre-symptomatic stage
Control
Pvalb-cKO
..25-12-01-D8_.0002.fcs
..12-01-D8_.0002.fcs\P1
..-01-D8_.0002.fcs\...\P2
P1
74.87%-
P1\P2
51.22%-T
P1\P2\P3
0.92%-T
..25-12-01-E8_.0002.fcs
..12-01-E8_.0002.fcs\P1
..12-01-E8_.0002.fcs\P1
P1
75.21%-
P1\P2
27.85%-T
P1\P3
12.04%-T
SSC-A
FSC-A
APC-A
CD64
TLR4
B)
Post-symptomatic stage
Page 1 / 2
..25-11-12-B7_.0002.fcs
..11-12-B7_.0002.fcs\P1
..-12-B7_.0002.fcs\...\P2
P1\P2
45.61%-T
P1\P2\P3
1.40%-T
..25-11-12-C7_.0002.fcs
..11-12-C7_.0002.fcs\P1
..-12-C7_.0002.fcs\...\P2
P1
90.79%-T
P1\P2
46.84%-T
P1\P2\P3
14.16%-T

**Figure S4**

**Figure S4: Flow cytometry analysis of TLR4 and CD64-positive cell populations during disease progression.**

(A) Representative flow cytometry gating strategy and analysis at the pre-symptomatic stage in control and *Pvalb*-cKO samples. Cells were first gated based on forward scatter (FSC-A) and side scatter (SSC-A) to identify the main cell population (P1). Cells were subsequently gated based on fluorescence intensity. Representative plots show CD64 versus TLR4 expression, indicating the proportion of $CD64^{+}TLR4^{+}$ cells. **(B)** Flow cytometry analysis at the post-symptomatic stage using the same gating strategy. Cells were gated by FSC-A/SSC-A, followed by fluorescence-based gating. Representative plots show CD64 versus TLR4 expression.

A)

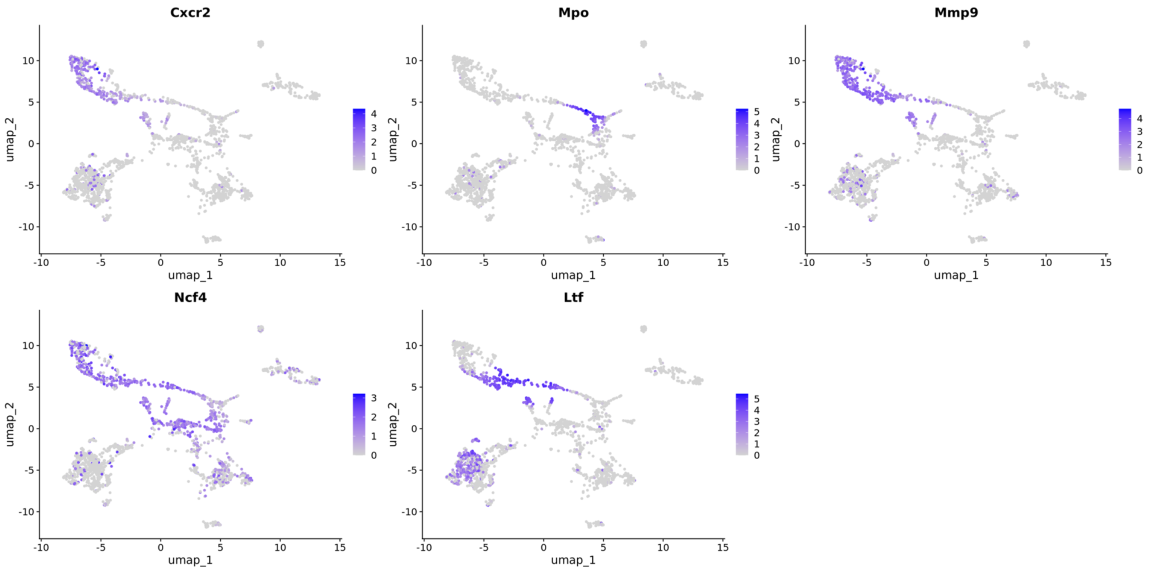
Cxcr2
Mpo
Mmp9
Ncf4
Ltf
umap_1
umap_2

B)

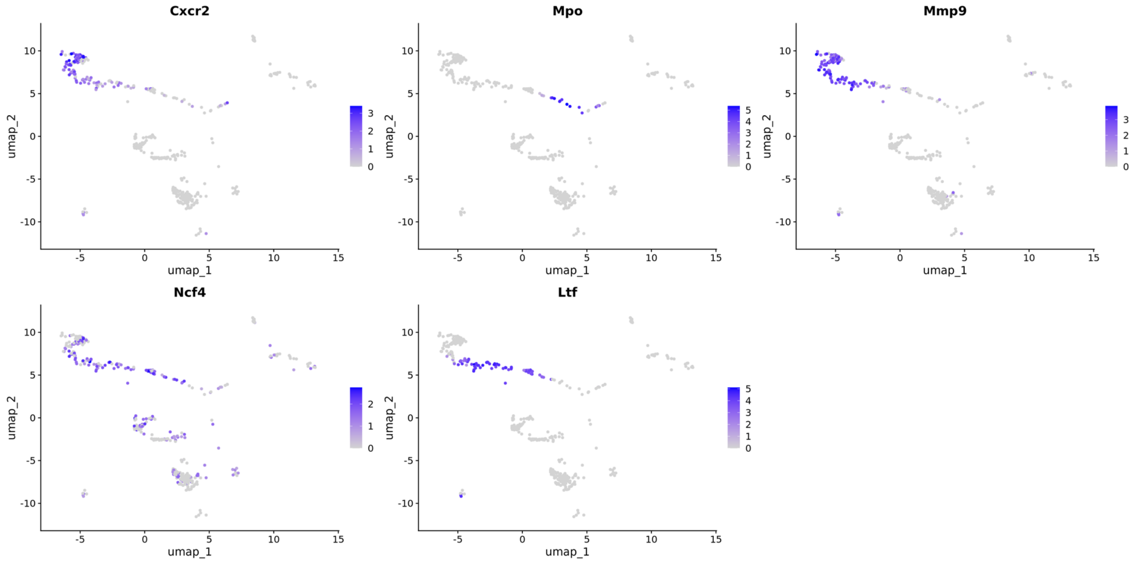
Cxcr2
Mpo
Mmp9
Ncf4
Ltf
umap_1
umap_2

### Figure S5

**Figure S5: UMAP visualization of neutrophil-associated gene expression across cellular populations.**

**(A)** Feature plots showing the expression of selected genes across the integrated UMAP embedding of single-cell transcriptomic data, including *Cxcr2, Mpo, Mmp9, Ncf4*, and *Ltf*. **(B)** UMAP feature plots highlighting the distribution of the same genes (*Cxcr2, Mpo, Mmp9, Ncf4*, and *Ltf*) within the macrophage cluster. Color intensity (blue gradient) represents relative gene expression levels, with darker shades indicating higher expression. Gray dots represent cells with low or undetectable expression.

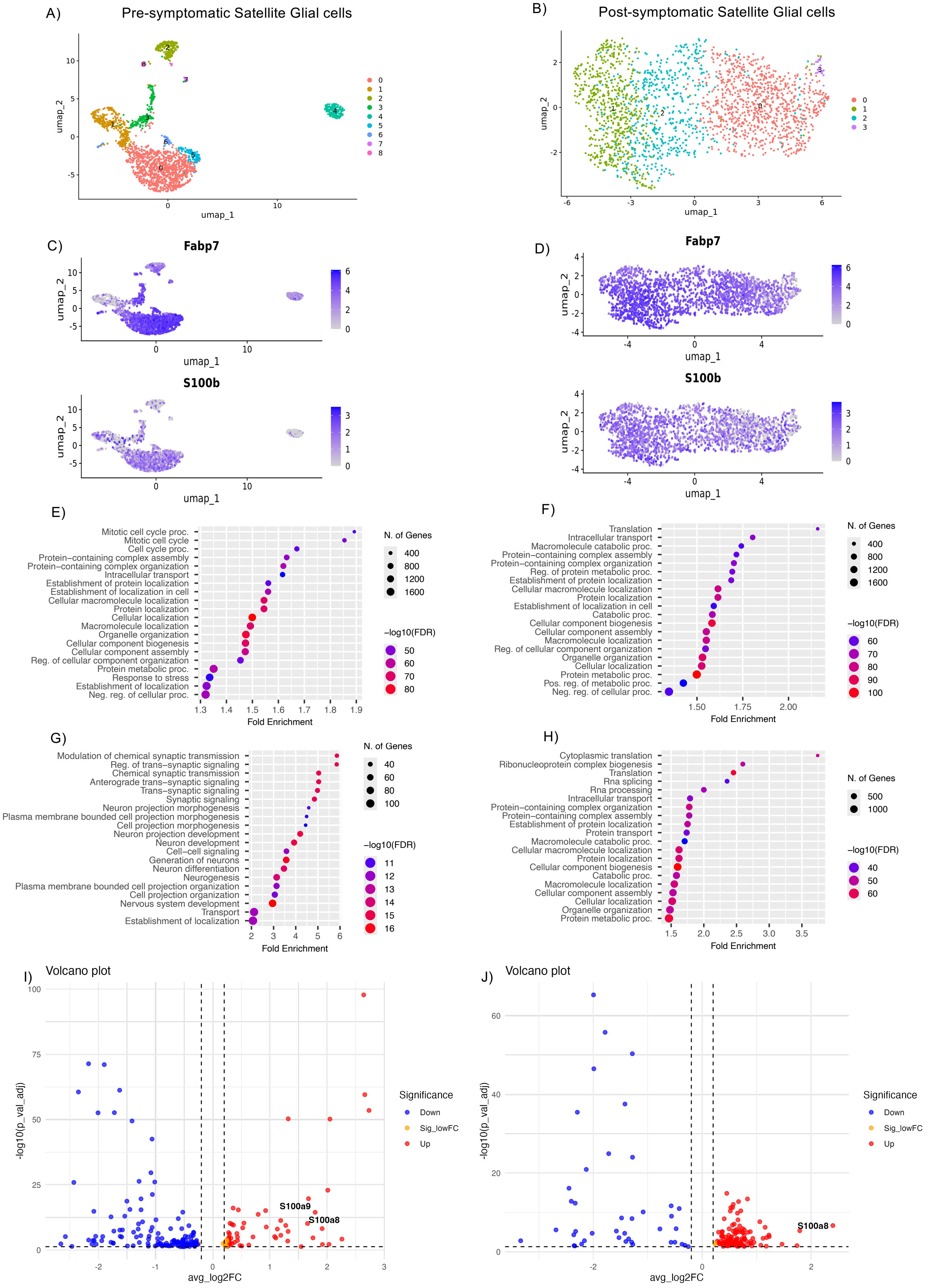
A)
Pre-symptomatic Satellite Glial cells
umap_1
umap_2
B)
Post-symptomatic Satellite Glial cells
umap_1
umap_2
C)
Fabp7
S100b
umap_1
umap_2
D)
Fabp7
S100b
umap_1
umap_2
E)
Mitotic cell cycle proc.
Mitotic cell cycle
Cell cycle proc.
Protein−containing complex assembly
Protein−containing complex organization
Intracellular transport
Establishment of protein localization
Establishment of localization in cell
Cellular macromolecule localization
Protein localization
Cellular localization
Macromolecule localization
Organelle organization
Cellular component biogenesis
Cellular component assembly
Reg. of cellular component organization
Protein metabolic proc.
Response to stress
Establishment of localization
Neg. reg. of cellular proc.
Fold Enrichment
N. of Genes
−log10(FDR)
F)
Translation
Intracellular transport
Macromolecule catabolic proc.
Protein−containing complex assembly
Protein−containing complex organization
Reg. of protein metabolic proc.
Establishment of protein localization
Cellular macromolecule localization
Protein localization
Establishment of localization in cell
Catabolic proc.
Cellular component biogenesis
Cellular component assembly
Macromolecule localization
Reg. of cellular component organization
Organelle organization
Cellular localization
Protein metabolic proc.
Pos. reg. of metabolic proc.
Neg. reg. of cellular proc.
Fold Enrichment
N. of Genes
−log10(FDR)
G)
Modulation of chemical synaptic transmission
Reg. of trans−synaptic signaling
Chemical synaptic transmission
Anterograde trans−synaptic signaling
Trans−synaptic signaling
Synaptic signaling
Neuron projection morphogenesis
Plasma membrane bounded cell projection morphogenesis
Cell projection morphogenesis
Neuron projection development
Neuron development
Cell−cell signaling
Generation of neurons
Neuron differentiation
Neurogenesis
Plasma membrane bounded cell projection organization
Cell projection organization
Nervous system development
Transport
Establishment of localization
Fold Enrichment
N. of Genes
−log10(FDR)
H)
Cytoplasmic translation
Ribonucleoprotein complex biogenesis
Translation
Rna splicing
Rna processing
Intracellular transport
Protein−containing complex organization
Protein−containing complex assembly
Establishment of protein localization
Protein transport
Macromolecule catabolic proc.
Cellular macromolecule localization
Protein localization
Cellular component biogenesis
Catabolic proc.
Macromolecule localization
Cellular component assembly
Cellular localization
Organelle organization
Protein metabolic proc.
Fold Enrichment
N. of Genes
−log10(FDR)
I)
Volcano plot
S100a9
S100a8
avg_log2FC
-log10(p_val_adj)
Significance
Down
Sig_lowFC
Up
J)
Volcano plot
S100a8
avg_log2FC
-log10(p_val_adj)
Significance
Down
Sig_lowFC
Up

**Figure S6**

**Figure S6: Single-cell transcriptomic characterization of satellite glial cells during disease progression.**

**(A)** UMAP visualization of satellite glial cells (SGCs) identified at the pre-symptomatic stage, showing transcriptionally define clusters (0–8).**(B)** UMAP visualization of SGC populations at the post-symptomatic stage, showing transcriptionally defined clusters (0–3). **(C–D)** Feature plots showing expression of SGC marker genes *Fabp7* and *S100b* across UMAP embeddings in the pre-symptomatic **(C)** and post-symptomatic **(D)** stages. Color intensity (blue gradient) represents relative gene expression levels, with darker shades indicating higher expression. **(E-F)** Gene Ontology (GO) biological process enrichment analysis of genes associated with SGC clusters, at pre-symptomatic (**E**) and post-symptomatic (**F**) stages. Dot size represents the number of genes associated with each term, and color indicates statistical significance (–log10 FDR). **(G-H)** Gene Ontology (GO) functional enrichment analysis of genes associated with SGC clsters at pre-symptomatic (**G**) post-symptomatic (**H**) stages. **(I–J)** Volcano plots showing differential gene expression in SGCs at pre-symptomatic **(I)** and post-symptomatic **(J)** stages. The x-axis represents the average log2 fold change (avg_log2FC) and the y-axis shows –log10(adjusted p-value). Red points indicate upregulated genes, blue points represent downregulated genes, and gray points indicate non-significant genes. Selected genes are labeled.
